# Cooperative and non-cooperative behaviour in the exploitation of a common renewable resource with environmental stochasticity

**DOI:** 10.1101/2020.01.09.901025

**Authors:** Michael Hackney, Alex James, Michael J. Plank

## Abstract

Classical fisheries biology aims to optimise fisheries-level outcomes, such as yield or profit, by controlling the fishing effort. This can be adjusted to allow for the effects of environmental stochasticity, or noise, in the population dynamics. However, when multiple fishing entities, which could represent countries, commercial organisations, or individual vessels, can autonomously determine their own fishing effort, the the optimal action for one fishing entity depends on the actions of others. Coupled with noise in the population dynamics, and with decisions about fishing effort made repeatedly, this becomes an iterated stochastic game. We tackle this problem using the tools of stochastic optimisation, first for the monopolist’s problem and then for the duopolist’s problem. In each case, we derive optimal policies that specify the best level of fishing effort for a given stock biomass. Under these optimal policies, we can calculate the equilibrium stock biomass, the expected long-term return from fishing and the probability of stock collapse. We also show that there is a threshold stock biomass below which it is optimal to stop fishing until the stock recovers. We then develop an agent-based model to test the effectiveness of simple strategies for responding to deviations by an opponent from a cooperative fishing level. Our results show that the economic value of the fishery to a monopolist, or to a consortium of fishing agents, is robust to a certain level of noise. However, without the means of making agreements about fishing effort, even low levels of noise make sustained cooperation between autonomous fishing agents difficult.

## 1. Introduction

Fish stocks are noisy [1, 2]. Sources of noise include: seasonality and variations in climatic conditions [3, 4]; interspecific interactions [5]; variations in nutrient availability [6]; spatial heterogeneity [7, 8]; differences in individual characteristics [9, 10]; and individual fecundity [11, 12]. Noise can affect population sizes continuously throughout the growing season, and in discrete episodes such as spawning events. Variability in recruitment is one the largest sources of noise [1, 6, 13].

A Markov decision process is a stochastic optimisation tool that couples a stochastic process with a series of decisions and associated payoffs [14]. A solution to a Markov decision process is a policy, which specifies the action to take as a function of the current state of the process, that maximises the total expected payoff over the lifetime of the process [15, 16]. Markov decision processes have been used to model optimal fishing efforts in the presence of environmental noise [17, 18]. These have been applied to age- and sex-structured populations [19, 20], multi-species fisheries [21], and in situations with incomplete information [22].

Classical fisheries biology is concerned with finding the fishing mortality rate that optimises fishery-level outcomes [23]. This applies in a situation where fishing effort is centrally controlled by a single regulator or a consortium of all relevant stakeholders. However, it does not apply in cases where multiple fishing entities independently set their own fishing efforts to meet their own objectives [24]. Fishing entities could be any unit with access to the fishery that has autonomy over its own fishing effort, and could represent countries, commercial organisations, or vessels. Decision-making by multiple fishing entities harvesting a shared stock is a game-theoretic problem because the state of the stock, and therefore each entity’s catch, is affected by the actions of the other fishers.

McKelvey [25] contextualised multinational fisheries management as involving brief and intense fish wars characterised by withdrawals from cooperative agreements, economic downturns in fishing communities, and heavily depleted fish stocks. Non-cooperative game theory offers a route to understanding such conflicts. Game theory has been used to investigate decisions about fishing effort by two countries exploiting a shared or partially shared fish stock [25–28], and to evaluate the allocation of spatio-temporal allocation of fishing [24, 29]. Martin-Herran and Rincón-Zapatero [30] and Chiarella et al. [28] found conditions under which games representing fisheries have Nash equilibria that are Pareto efficient. This is of interest because Pareto efficient solutions are socially optimal and it is unusual for Nash equilibria to have this property. Game theoretic concepts have also been used in models of small-scale fisheries with a population of autonomous fishing agents [31, 32]. These studies showed that an approximate Nash equilibrium emerged among fishing agents and produced a pattern of fishing selectivity that was well matched with the natural productivity of ecosystem components. However, the open-access case, where there is no prescribed limit to fishing effort, resulted in overfishing. The emergence of a Nash equilibrium among fishing agents is analogous to the ideal free distribution of foraging theory [33, 34], which predicts that that foraging effort is spatially distributed in proportion to resource productivity [35, 36]. This concept has also been applied to fleet dynamics to quantify the spatial distribution of fishing effort [37–39].

Most modelling studies that take a game theoretic approach to decisions about fishing effort do not explicitly account for the effects of noise in the fish stock. McKelvey [25] modelled an interception fishery, where the stock is harvested sequentially by two competing countries as it migrates through their exclusive economic zones, but only included stochasticity in the payoff function, not in the stock itself. Sobel [40] considered a stochastic game model of a fishery, which required that the probability of the process transitioning to a given state could only depend on the current action and not the current state. This is unrealistic for most fishing scenarios, where the spawning stock biomass in one year is a significant determinant of biomass in subsequent years. The absence of noise from most game theoretic models is an important limitation because noise not only has a major effect on stock biomass, and therefore the likelihood of overfishing, but also on the information available to fishing agents when deciding on their fishing effort.

In this paper, we present a new method for calculating optimal fishing effort, both for a monopolist and a duopolist, in noisy conditions. A fishing agent’s optimal effort in any given period is the effort that maximises that agent’s total expected future net profits, given the current stock biomass. First, we formulate the monopolist’s problem as a Markov decision process [14]. The state variable, representing stock biomass, and the action variable, representing fishing effort, are both continuous. A solution to this process is a policy that specifies the optimal fishing effort for any level of stock biomass.

Secondly, we extend our analysis to the case of a duopolist by reformulating the model as a stochastic Markov game [41]. A Nash equilibrium solution of the Markov game is a solution in which, in any given period, both players are following a policy that is a best response to the other player’s policy. Although finite Markov decision processes have a (possibly non-unique) stationary optimal policy [14], this is not necessarily the case for Markov games [42, 43]. Most work on stochastic games has focused on the zero-sum case, which does have a unique Nash equilibrium [42], whereas the problem considered here gives rise to a non-zero-sum game. Methods used to solve non-zero-sum stochastic games include value iteration [44], policy iteration [41, 45, 46], and reinforcement learning [47, 48]. Reinforcement learning is particularly useful for problems with a high-dimensional state space and/or an unknown model structure. Reinforcement learning algorithms often use a mixture of exploitation (taking the action believed to give the best long-term return with current information) and exploration (taking a random action in order to collect information on the problem structure) [49]. These techniques have been studied in various applications, including operations research and control theory [50], robotics [51], neuroscience [52], and finance [53], but not to our knowledge exploitation of a shared renewable resource.

The problem analysed in this paper has a sufficiently low-dimensional state space that we can use a direct optimization method based on value iteration. This method converges to a solution known as a stationary open-loop equilibrium, meaning that players’ actions depend on the current state of the process and the anticipated current and future actions of the other player, but not the previous actions of the other player [54]. Open-loop equilibria are analytically tractable, but restrict the type of strategies that fishing agents may use. We therefore formulate an agent-based model to evaluate the effectiveness of simple closed-loop strategies, in which players can modify their actions based on their beliefs about their opponent’s previous actions [27].

We show that the decision problem faced by two autonomous agents (i.e. duopolists) that cannot make binding agreements has the structure of prisoner’s dilemma. We show that the presence of noise in the fish stock drastically reduces the potential for cooperation between duopolists. Intuitively, this occurs because, in years when the biomass is lower than expected, it is impossible for a duopolist to know whether this is caused by noise or by “cheating” (i.e. higher than mutually optimal levels of fishing effort) by their opponent.

## 2. Markov decision process model

### 2.1. Stock biomass model

The stock biomass model consists of two components: an ecological component representing the change in population biomass due to natural mortality and recruitment from one year to the next; a harvesting component representing reduction in biomass due to fishing. Generational spawning is common in fish species, typically occurring annually [55–57], over short spawning seasons [58]. We therefore assume that recruitment occurs as discrete, annual events, whereas harvesting occurs continuously throughout the year.

Firstly, for the ecological component, we use a Beverton–Holt function plus a normally distributed noise term. The Beverton–Holt function models the dimensionless biomass of new recruits *f*(*x*) in a given year as a function of the spawning stock biomass *x* in the previous year:

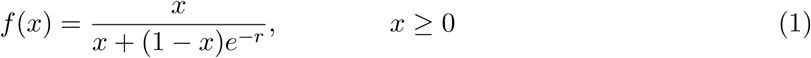

where *r* is the intrinsic growth rate (per year). This is the discrete-time version of the logistic growth differential equation and is a classical stock–recruitment relationship in fisheries biology, modelling density-dependent adult spawning and larval survival [59]. The function in Eq. (1) is a saturating function of biomass with a unique stable equilibrium at *x* = 1, which corresponds to the carrying capacity of the system. Alternative stock–recruitment relationships exist, notably the Ricker [60] model which assumes recruitment decreases when stock biomass exceeds a threshold level. However, for simplicity we do not investigate these here.

We include in the model a noise term, representing stochasticity in the recruitment process, which is assumed to be normally distributed and independent from one year to the next. Combining this with the Beverton-Holt map, the biomass after recruitment in year *n* is given by

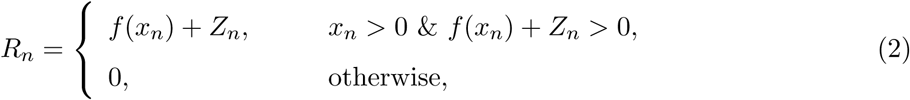

where *x*_*n*_ is the biomass at the beginning of year *n* and the *Z*_*n*_ are independent, identically distributed normal random variables *N* (0, *σ*^2^). The parameter *σ* represents the magnitude of the noise in the ecological dynamics. Note that if the biomass ever becomes non-positive, it is assumed to remain at zero for all future time representing extirpation of the stock, i.e. zero is an absorbing state of the stochastic process.

For the harvesting component of the model, we assume that in year *n* the total fishing mortality rate *F*_*n*_ (per year) is constant throughout the year. For simplicity we assume that all other sources of mortality are accounted for by the map in Eq. (2). This means that the biomass at the beginning year *n* + 1 is

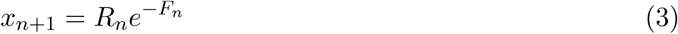

and the total yield extracted during year *n* is

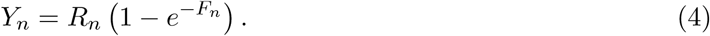

The harvesting component is assumed to be deterministic, i.e. there is no noise in either the fishing mortality rate *F*_*n*_ or the yield *Y*_*n*_.

Combining Eqs. (2) and (3) gives a stochastic mapping from the biomass at the beginning of one year to the biomass at beginning of the next year:

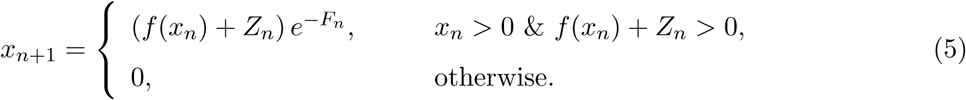

According to Eq. (5), if the biomass at the beginning of year *n* is *x* and fishing mortality rate in year *n* is *F*, then the probability density of having biomass *y* > 0 in year *n* + 1 is

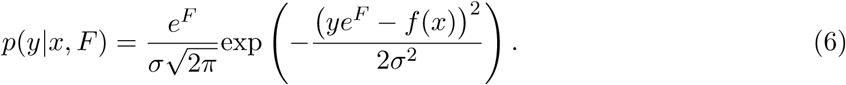

The probability of having zero biomass in year *n* + 1 is given by 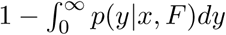.

Figure 1 illustrates the sequence of biomass observation, recruitment, noise and fishing. It is possible to model this sequence in two different ways, either with recruitment and noise occurring before fishing (Fig. 1a) or vice versa (Fig. 1b). Under the first model, fishing agents must decide on their level of fishing effort before the stock biomass is affected by noise. Under the second model, fishing agents can effectively observe the sign and magnitude of that year’s noise term before having to decide their fishing effort. This makes a small difference to the optimal policies that the model produces because agents can better tune their effort levels to current environmental conditions. However, in the duopolist case (see Sec. 2.3), having knowledge of the stock biomass immediately before the fishing season begins means that each fishing agent can infer, with perfect accuracy, the actions of their competitor via their own yield. To retain some realistic uncertainty in beliefs about other agent’s actions, we therefore use the sequence shown in Fig. 1a. This means that agents cannot observe the noise process directly and hence cannot distinguish the effects of noise and competitor actions.

**Figure 1:**
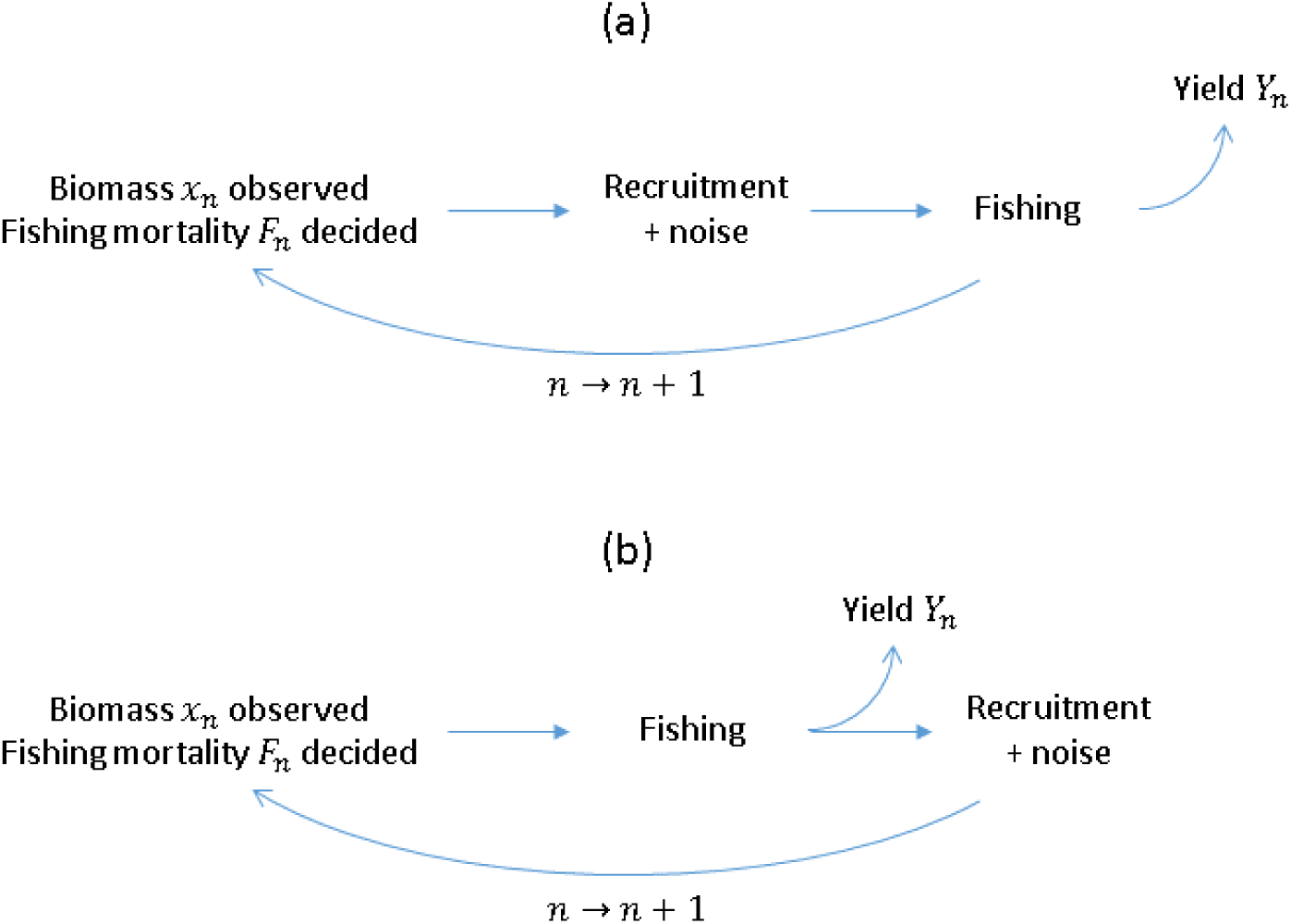
Schematic diagram showing two possible ways of modelling the annual sequence of biomass observation, recruitment and noise, and fishing. In (a), a fishing agent cannot distinguish the effects of noise from fishing by other agent(s). In (b), a fishing agent can infer perfectly the total fishing mortality applied by the other agent(s) because the combination of the biomass observation *x*_*n*_ at the start of year *n* and the yield *Y*_*n*_ in year *n* allow them to separate the effects of noise and fishing mortality. To avoid fishing agents having perfect information, we model the sequence as shown in (a).

### 2.2. Monopolist problem

We first consider the optimisation problem faced by a single agent with the ability to observe the biomass *x*_*n*_ at the beginning of each year and subsequently determine the total fishing mortality *F*_*n*_ to apply that year. This is a Markov decision process with a continuous state space *S* = {*x* ∈ ℝ : *x* ≥ 0} and continuous action space *A* = {*F* ∈ ℝ : *F* ≥ 0}.

We assume that biomass extracted from the fishery has a unit price *b* so that the revenue in year *n* is *bY*_*n*_, where *Y*_*n*_ is the yield in year *n* as defined in Eq. (4). Similarly, we assume that there is a cost *c* per unit fishing effort so that the expenditure in year *n* is *cF*_*n*_. By rescaling the profit relative to the unit price *b*, we may set *b* = 1 without less of generality. Hence, if the biomass at the beginning of the year is *x*, the fishing mortality rate is *F*, and the biomass at the beginning of the following year is *y* then the net profit *P* is

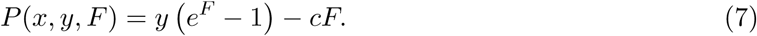

For generality, the profit function *P* is a function of the current state of the process (*x*), the action taken (*F*), and the subsequent state to which the process transitions (*y*). In this case, because of the sequence shown in Fig. 1a, the profit is a deterministic function of the biomass at the end of the fishing period, i.e. the subsequent state of the process *y*, independent of the current state *x*.

For a time horizon of *N* years, the monopolist’s problem is to choose a policy *Q*_*n*_(*x*), which specifies the fishing mortality *F* to apply when the biomass is state *x* in year *n* (*n* = 1, …, *N*). An optimal policy is one that maximises the expected value of the total discounted profit

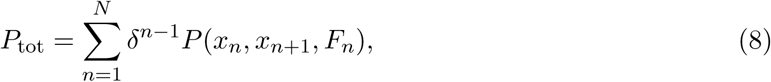

where *δ* ∈ [0, 1] is the discount factor. This stochastic optimisation problem may be solved via the Bellman equation [61] for the expected future discounted profit *V*_*n*_(*x*), referred to as the value, when the biomass is *x* in year *n*. The Bellman equation expresses the value recursively as the expected profit in year *n*, plus the expected value in year *n* + 1, maximised over all possible actions in year *n*:

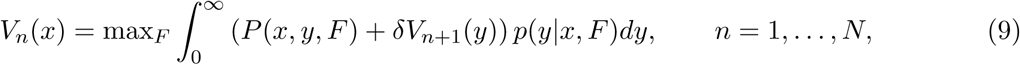

The value *V*_*N*+1_(*x*) in year *N* + 1 is defined to be zero for all states *x*. This means that year *N* is the last year in which any revenue may be extracted from the resource. Note that the value *V*_*n*_(0) of being in state *x* = 0 is zero for all *n*. This is because, if the process is in state *x* = 0 in year *n*, it will remain in state *x* = 0 for all future years with probability 1, regardless of the actions taken. Hence no current or future revenue can possibly be extracted. The Bellman equation reduces a decision problem into smaller subproblems according to Bellman’s principle of optimality, which states that “an optimal policy has the property that whatever the initial state and initial decision are, the remaining decisions must constitute an optimal policy with regard to the state resulting from the first decision” [62].

We solve the Bellman equation by a method referred to as value iteration [14] as follows. Eq. (9) with *n* = *N* and for a given value of *x* is a deterministic optimisation problem over the one-dimensional action variable *F*. This allows calculation of the optimal policy, denoted 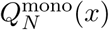, and the value function *V*_*N*_ (*x*) for year *N*. Once *V*_*N*_ (*x*) is known, Eq. (9) with *n* = *N* − 1 can be solved to find 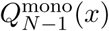 and *V*_*N*−1_(*x*), and so on. Hence, Eq. (9) can be solved by backward induction in *n* to find the optimal policy 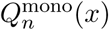 for *n* = 1, …, *N*.

If the optimal policy converges such that 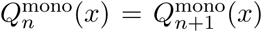, it is referred to as a stationary optimal policy. For problems where the one-stage profit *P* (*x, y, F*) is bounded, a stationary optimal policy is guaranteed to exist, though may not be unique [14]. Technically, in our model, *P* (*x, y, F*) is unbounded as the state variable *y* is unbounded. However, it is vanishingly unlikely for *y* to take values much larger than the carrying capacity *K*, so we expect a stationary optimal policy to exist in practice. If a stationary optimal policy exists, we seek a pseudo-stationary state distribution *π*(*x*), which we define as a distribution satisfying

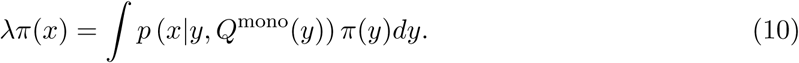

If the process is in a pseudo-stationary distribution *π*(*x*) and the monopolist is following a stationary optimal policy, then the process has probability 1 − λ per year of entering the absorbing state *x* = 0 (i.e. the stock is extirpated). Conditional on non-extinction, the process will remain in the pseudo-stationary distribution. If the value function also converges such that *V*_*n*_(*x*) = *V*_*n*+1_(*x*), we can also calculate the expected long-term discounted return from fishing *V*. This is given by the expected value of *V* (*x*) when the process is in a pseudo-stationary distribution:

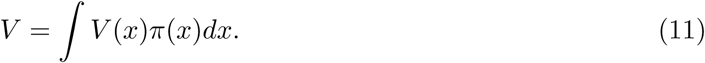

### 2.3. Duopolist problem

We now consider the situation in which two agents must independently decide the fishing mortality to apply each year. The total fishing mortality in year *n* is 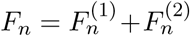, where 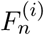 is the fishing mortality applied by agent *i* ∈ {1, 2} in year *n*. In this case, the total yield is still determined by Eq. (4), but this is now split between the two agents according to their relative fishing mortality rates. The agents may have different cost to price ratios *c*_*i*_ and discount factors *δ*_*i*_. If the biomass at the beginning of the year is *x*, agent *i* applies fishing mortality *F* ^(*i*)^, and the biomass at the beginning of the following year is *y*, then the net profit *P* ^(*i*)^ for agent *i* is

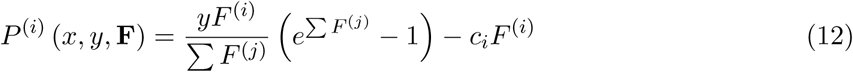

where **F** = [*F* ^(1)^, *F* ^(2)^] and all summations are over *j* = 1, 2.

The duopolist’s problem can also be formulated as a Bellman equation, conditional on knowing what action the other agent will take when the process is in a given state. Suppose that agent *j* applies fishing mortality *F* ^(*j*)^. Then the Bellman equation for agent *i* in year *n* is:

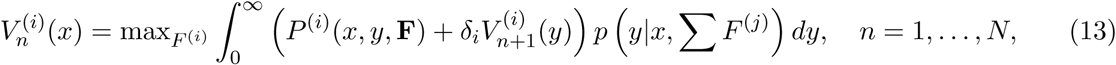

The optimal fishing mortality for agent *i* when the process is in state *x* will depend on the fishing mortality applied by *j*. We call a pair of policies 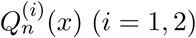 in year *n* optimal policies if they are best responses to each other, i.e. 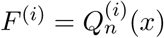 is the solution of Eq. (13) when 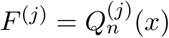.

In the symmetric case where the agents are identical (*c*_1_ = *c*_2_, *δ*_1_ = *δ*_2_), there must be a pair of optimal policies that are the same for both agents, i.e. 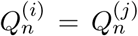. This constraint can be used to to reduce Eq. (13) to a single-variable optimisation problem in each year (see Appendix A for details). As with the monopolist problem, the optimal policies are found via backward iteration in *n*. We denote the symmetric duopolist’s optimal policy in year *n* by 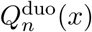. If the duopolist’s optimal policy converges to a stationary policy, we can calculate a pseudo-stationary distribution, the probability of extirpation, and the expected long-term return from fishing via equations analogous to Eq. (10) and (11). In the asymmetric case, optimal policies in year *n* are found by iteratively optimising agent *i*’s policy in response to a fixed policy by agent *j* and vice versa until convergence (see Appendix A for details).

### 2.4. Results

Figure 2 shows the monopolist’s and symmetric duopolist’s optimal policy and value function for a *N* = 40 year period when *σ* = 0.1. This noise level means that the standard deviation of annual noise is 10% of the population carrying capacity. The monopolist’s optimal policy in the final year (yellow curve in Fig. 2a) maximises net profit in a single, fixed time period without regard to the biomass remaining at the end of this period. As expected, the optimal policy is always an increasing function of biomass, so it is optimal to fish harder when the stock biomass is high. As the process iterates backwards in time, the optimal policy quickly converges to a stationary policy: the policies in all years except the final year (black curve in Fig. 2a) are almost identical. In any given year, there is a threshold biomass below which it is optimal to cease fishing because the cost of fishing would exceed the revenue generated. In the final year (*n* = *N*), the optimal fishing mortality is higher, and the threshold biomass for fishing is lower, than in preceding years (*n* ≤ *N* − 1). This is because in the final year there is no incentive to protect the stock for future periods. The value function increases as the process iterates backwards in time (darker curves in Fig. 2b) and eventually converges towards a stationary value, but this takes longer than convergence of the optimal policy.

**Figure 2:**
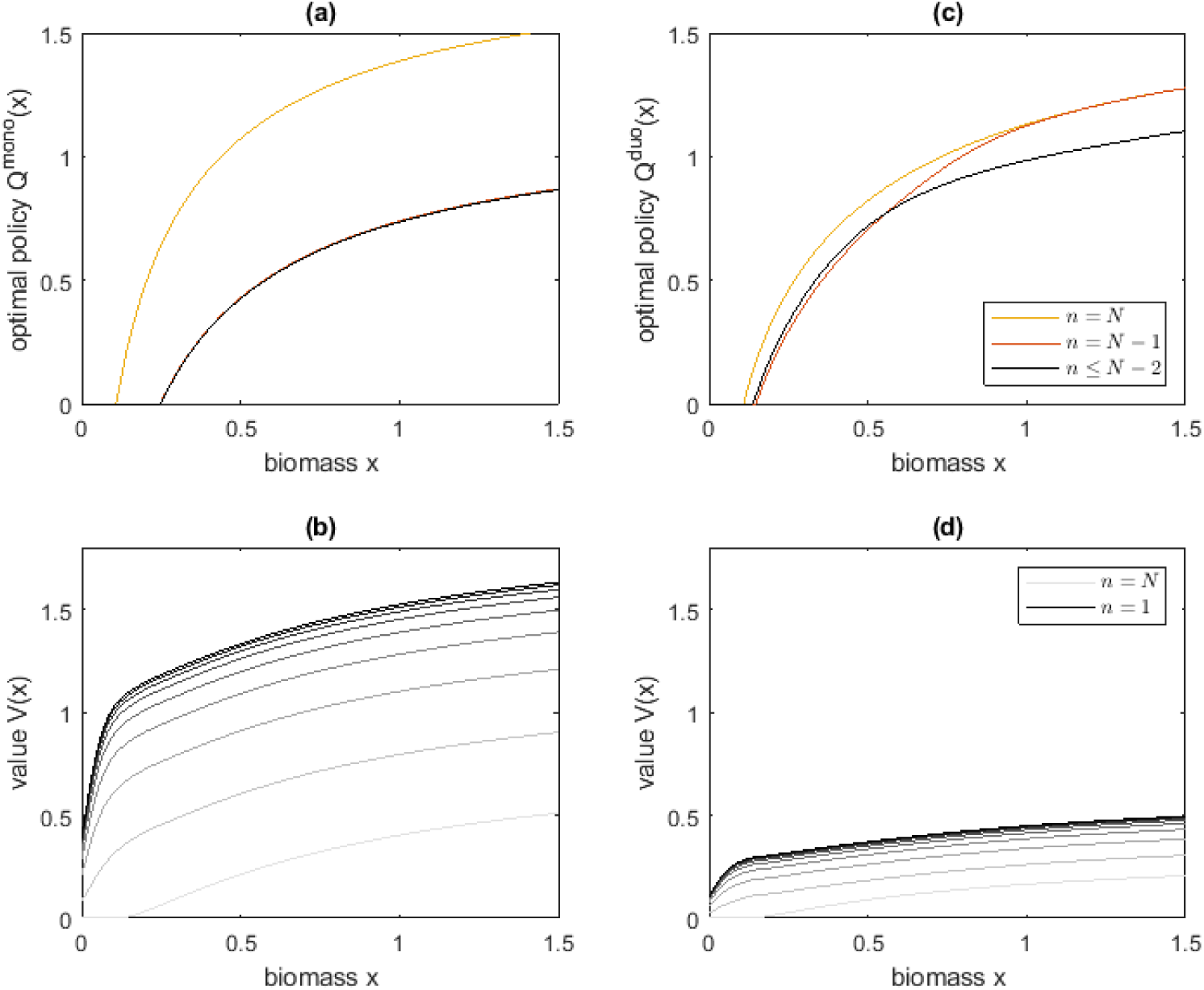
Solutions to the Markov decision process over a *N* = 40 year period for: (a,b) a monopolist; and (c,d) a symmetric duopolist. (a,c) show the optimal policy in the final year (yellow), penultimate year (red), and preceding years (black) for which the optimal policy has converged to a stationary policy in both cases. (b,d) show the value function in the final year *n* = *N* (lightest shade) through to the first year *n* = 1 (darkest shade) at 5 year intervals. Noise level *σ* = 0.1. Other parameter values shown in Table 1

The symmetric duopolist’s optimal policy (Fig.2c) follows a similar shape to the monopolist’s but, in all except the final year, specifies a higher fishing mortality for a given biomass *x*, and has a lower threshold biomass required for fishing. The value function for a symmetric duopolist (Fig.2d) is less than half that of the monopolist, indicating that the economic performance of the fishery as a whole is reduced when exploited by two non-cooperative agents. The duopolist’s optimal policy converges to a stationary policy within 2 years of the final time period and, as in the monopolist case, the value function converges but more slowly. For the remainder of the paper, we focus on the stationary solution to the Markov decision process (i.e. long time horizon) found by solving the Bellman equation over a *N* = 40 year period and examining the solution in year *n* = 1.

**Table 1:**
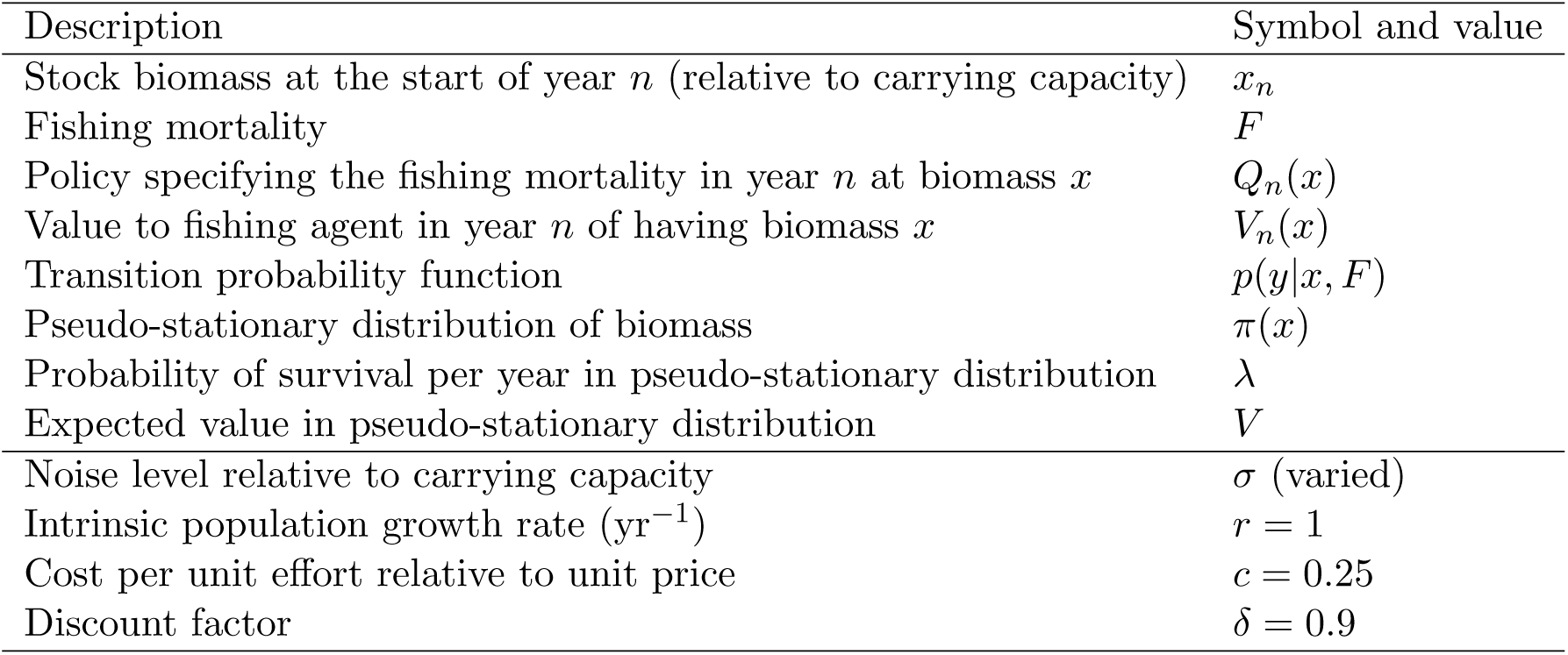
Model variables and parameter values.

Figure 3 shows the monopolist’s and symmetric duopolist’s stationary optimal policy and value function, and the pseudo-stationary biomass distribution, for a range of noise levels *σ*. When noise is small, the pseudo-stationary biomass distribution under a monopolist (Fig. 3c) is tightly peaked, meaning there is little variation from year to year. The mean of the biomass distribution is slightly below the equilibrium biomass at maximum economic yield (*B*_MEY_) under the deterministic version of the model (vertical dotted line). This reference point is calculated by maximising the net profit when the biomass is at equilibrium and there is no noise (*σ* = 0). The slight discrepancy is because the classical MEY solution does not discount future profits (although it can be adjusted to do so), so it is optimal to main stock biomass at a slightly higher level. The value of the resource to the monopolist with low noise (dark blue curve in Fig. 3b) also matches with the MEY prediction.

**Figure 3:**
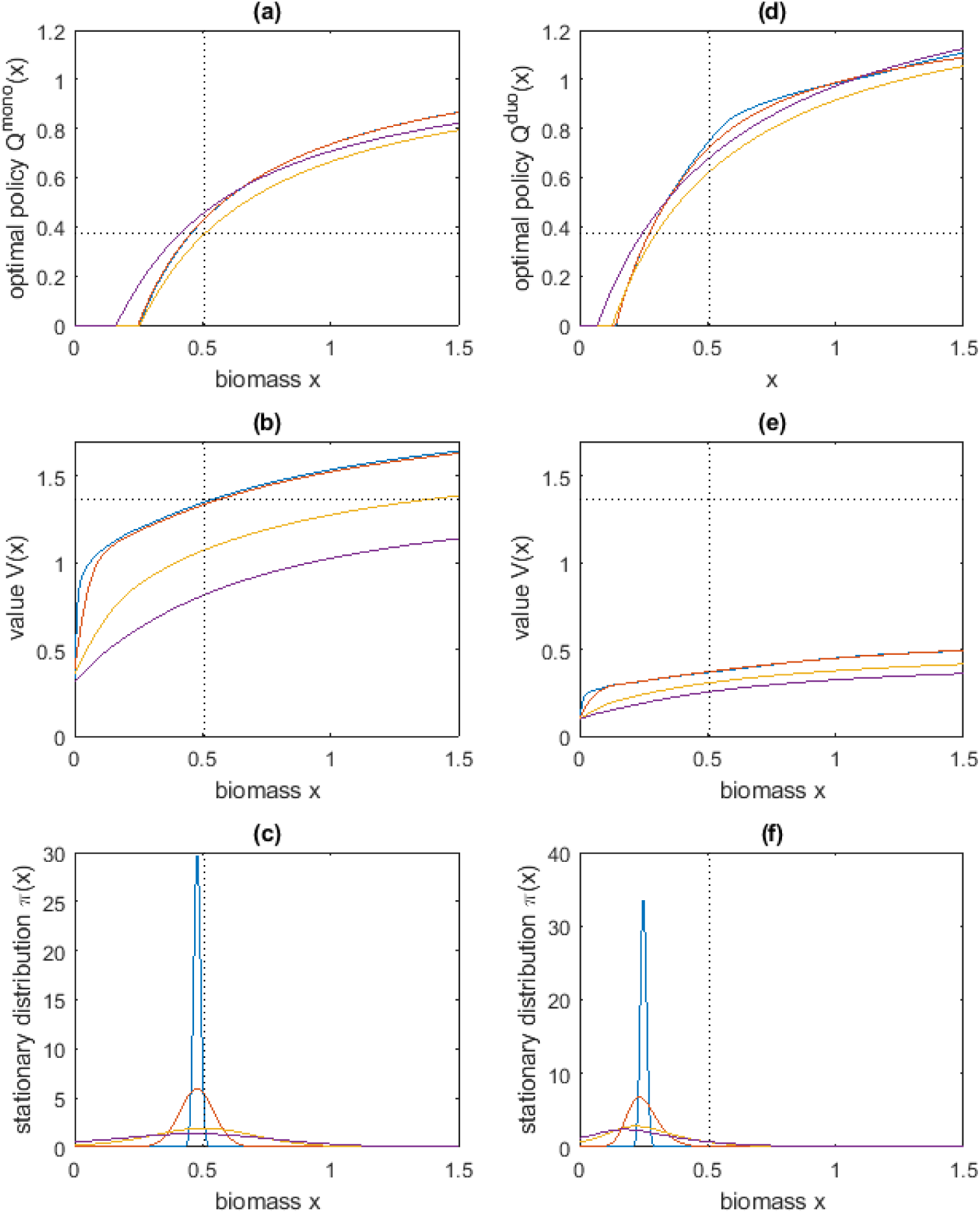
Effect of noise on the stationary optimal policy, value function and biomass distribution for: (a-c) a monopolist and (d-f) a symmetric duopolist. Noise levels *σ* = 0.02 (dark blue), 0.1 (red), 0.3 (yellow), 0.5 (purple). Panels (a,d) show the stationary optimal policy; (b,e) show the stationary value function; (c,f) show the pseudo-stationary biomass distribution. Dotted lines show the equilibrium biomass, fishing mortality and value corresponding to the maximum economic yield (MEY) in the deterministic monopolist case. Parameter values shown in Table 1

In the case of a symmetric duopoly, the optimal fishing mortality for each agent is higher than for a monopolist at the same stock biomass, and the threshold required for fishing is lower than for a monopolist (Fig. 3d). The mean of the pseudo-stationary biomass distribution (Fig. 3f) is approximately half that under a monopolist, and the value is less than half that of the monopolist (Fig. 3e). This shows that two non-cooperative fishing agents will deplete the stock below optimal levels and receive sub-optimal returns.

As noise increases, the shape of the optimal policy changes slightly and the threshold biomass required for fishing decreases (Fig. 3a,d). This reflects a detrimental consequence of noise on the bioeconomic dynamics. When noise is small, agents will avoid fishing in years when the biomass is low in order to protect future profits. However, when noise is larger, there is more uncertainty to future profits, so less of an incentive to protect them. Increasing noise also decreases the value function (Fig. 3b,e). This shows that higher profits in high biomass years do not offset the reduction in expected long-term profits due to the risk of extirpation of the stock.

Figure 4 shows the effect of asymmetry in relatives costs *c*_*i*_ or discount factors *δ*_*i*_ on the stationary optimal policies and value functions in a duopoly. When one agent has a higher cost to price ratio *c*_*i*_ (red curve in Fig. 4a), that agent requires a higher threshold biomass for fishing and fishes at a lower rate than the other agent for any given biomass. When one agent has a lower discount factor *δ*, meaning they place less value on profits in future years relative to the current year (red curves in Fig. 4c), they require a slightly lower threshold biomass for fishing, and fish at a higher rate when biomass is relatively low (but above threshold). In both these cases, the agent with the higher cost or the lower discount factor unsurprisingly derives less long-term expected value from the stock at any given level of biomass (red curves in Fig. 4b,d). More interesting is the impact of this on the first agent’s value relative to the symmetric case shown in Fig. 2d. Competing with an opponent with higher costs increases the value to the first agent (blue curve in Fig. 4b) because their opponent is fishing less. In contrast, competing with an opponent with a low discount factor reduces the value for both agents (Fig. 4d). This is because the short-sighted actions of the opponent result in overfishing and hence suboptimal returns from the fishery as a whole.

**Figure 4:**
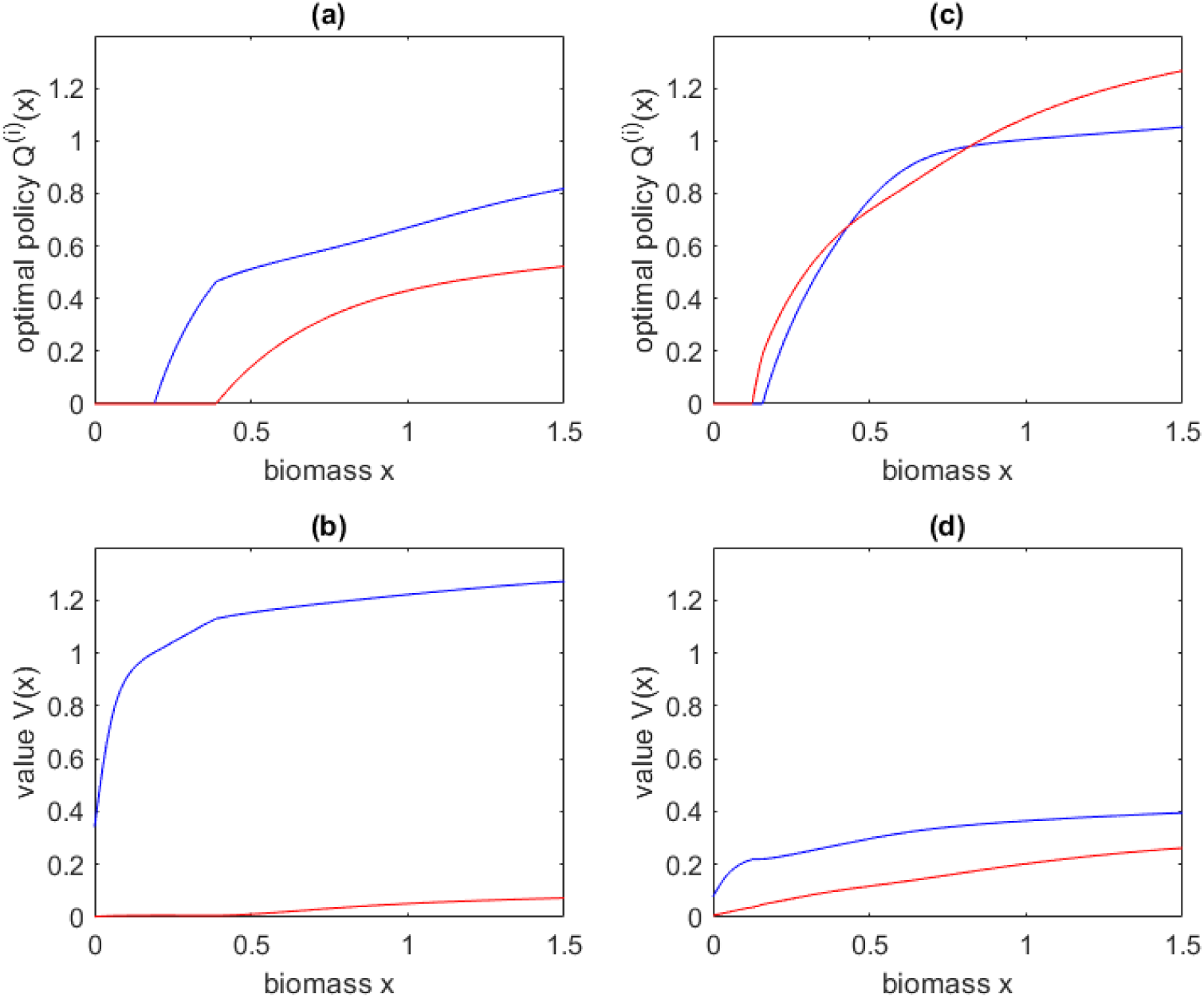
Effect of asymmetry in relative cost *c*_*i*_ or discount factor *δ*_*i*_ on the stationary duopolist’s: (a,b) optimal policies; and (c,d) and value functions. In (a,c), agent 1 (blue) has low relative cost and agent (red) has high relative cost (*c*_1_ = 0.25, *c*_2_ = 0.5). In (b,d), agent 1 (blue) has a high discount factor and agent 2 (red) has a low discount factor (*δ*_1_ = 0.9, *δ*_2_ = 0.5). Noise level *σ* = 0.1.

## 3. Closed-loop strategies

The stochastic optimisation framework developed in Sec. 2 allows for open-loop strategies, in which agents can base their choice of action on the current state of the process, and the inferred current and future actions of their competitor, but not on previous actions or states [54]. An open-loop strategy corresponds to a set of policies *Q*_*n*_ : *S* → *A*, that defines a unique action *F* ∈ *A* for each state *x* ∈ *S* in each year *n* = 1, …, *N*. This means that agents cannot directly respond to deviations by other agents from an equilibrium strategy. Closed-loop strategies allow an agent’s actions to depend in part on their beliefs about the past actions of other agents. In the context of our problem, a closed-loop strategy may be defined as a set of policies *Q*_*n*_ : *S*^*n*^ *× A*^*n*−1^ *×* ℝ^*n*−1^ → *A* that is a function of the history of states *x*_*k*_ and the agent’s actions *F*_*k*_ and one-stage profits *P*_*k*_ (*k* = 1, …, *n* − 1), as well as the current state *x*_*n*_. These strategies are not analytically tractable due the high dimensionality of the domain of *Q*_*n*_. Hence, in this section, we develop a stochastic agent-based model to investigate simple closed-loop strategies. We simulate the decision problem faced by a duopolist using an agent-based model with two agents.

### 3.1. Agent-based model

The optimal policy for the duopolist found by solving Eq. (13) is an open-loop Nash equilibrium. In other words, in a given year and for a given biomass, neither agent can increase their expected discounted profit, over the time horizon of the process, by unilaterally changing to a different fishing mortality. In contrast, a Pareto efficient strategy set is one where it is impossible to increase one agent’s expected return without decreasing the other agent’s expected return. In the symmetric duopoly model, any combination of agent fishing mortalities such that 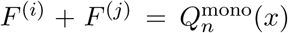 is Pareto efficient. For simplicity, we focus only on the symmetric Pareto efficient solution, i.e. 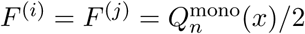.

The existence of a Nash equilibrium and a Pareto efficient solution means that the duopolist’s decision problem shares some characteristics of an iterated prisoner’s dilemma. In reality, agents can choose from a continuum of fishing mortalities *F* ^(*i*)^ ≥ 0. However, for simplicity, in this section we assume that each year they make a binary choice of either: (i) fishing according to the Pareto efficient solution 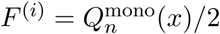; or (ii) fishing according the Nash equilibrium 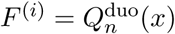. In keeping with the terminology of a prisoner’s dilemma, we label these two actions “cooperate” and “defect” respectively.

We compare three simple strategies: always cooperate (ALLC), always defect (ALLD), and tit-for-tat (TFT). In the classic iterated prisoner’s dilemma, each player can directly infer the action of their opponent in the previous round. TFT specifies that the player should cooperate in the first round and, in subsequent rounds, take the same action that their opponent took in the previous round [63]. This is known as a memory-one strategy as it requires the player to have memory of their opponent’s action one round previously. In our model, fishing agents cannot accurately infer the fishing mortality applied by the other agent in the previous round, because of noise. Nevertheless, agent *i* can calculate the anticipated yield in year *n* under the assumption their opponent cooperated, denoted 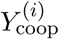, and under the assumption their opponent agent defected, denoted 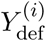:

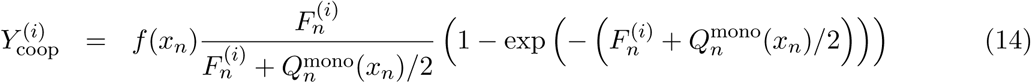

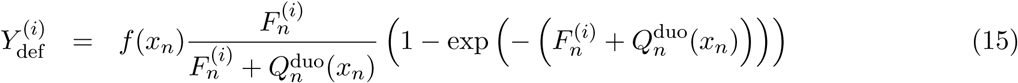

The agent then observes their actual yield 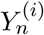 and we assume that they believe their opponent cooperated if 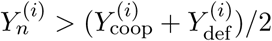, otherwise they believe their opponent defected.

For simplicity, we assume that the time-horizon is sufficiently long that the monopolist’s and duopolist’s optimal policies are stationary. Hence, for a given biomass *x*, the fishing mortalities corresponding to the cooperate action and the defect action are the same every year. A realisation of the agent-based model is simulated as follows:

1. Initialise the process by setting *n* = 1, *x*_*n*_ = *K* and *B*^(*i*)^ = 1 (*i* = 1, 2).
2. Both agents observe *x*_*n*_. Determine each agent’s fishing mortality *F* ^(*i*)^ according to *x*_*n*_ and the strategy they are following. An agent *i* playing TFT will cooperate if *B*^(*i*)^ = 1 and defect if *B*^(*i*)^ = 0.
3. Calculate *x*_*n*+1_ via the stochastic map in Eq. (5).
4. Calculate each agent’s yield 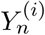 and net profit 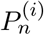 via Eq. (12).
5. For an agent playing TFT, calculate the anticipated yield under assumption the other agent cooperated 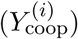, and under the assumption the other agent defected 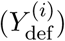 via Eqs. (14) and (15). Update agent’s belief by setting *B*^(*i*)^ = 1 if 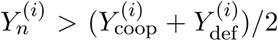 and *B*^(*i*)^ = 0 otherwise.
6. Increase *n* by 1 and repeat steps 2–6 until *n* = *n*_max_ or *x*_*n*_ = 0.

An initial burn-in period of *n*_0_ years is discarded. For each realisation, we calculate each agent’s total discounted profit,

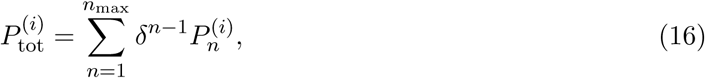

We also calculate the average probability of extirpation of the stock per year. These outputs are averaged over *M* independent realisations of the process. Any realisation in which the stock is extirpated before the end of the burn-in period is discarded.

### 3.2. Results

We simulated the agent-based model in three cases: both agents playing ALLC; both agents playing ALLD; and both agents playing TFT. Figure 5 compares agent-based model simulations with results from the Markov decision process from Sec. 2. With cooperative behaviour (ALLC, blue), the agent-based model agrees well with the Markov decision process under the monopolist’s optimal policy. With uncooperative behaviour (ALLD, red), simulations agree with the results of the duopolist’s Nash equilibrium. In both cases, noise levels below around 10% of carrying capacity do not appreciably affect the expected value (i.e. the average long-term return from fishing). This is because the probability of extirpation is negligible at these noise levels and, in the long run, the increased profit in high biomass years balances the reduced profit in low biomass years. However, increasing noise above around 10% of carrying capacity increases the probability of extirpation and therefore reduces the expected value. The probability of extirpation is higher under uncooperative behaviour than under cooperative behaviour, because the aggregate fishing mortality is higher and the equilibrium average biomass is lower, as shown by Fig. 3.

**Figure 5:**
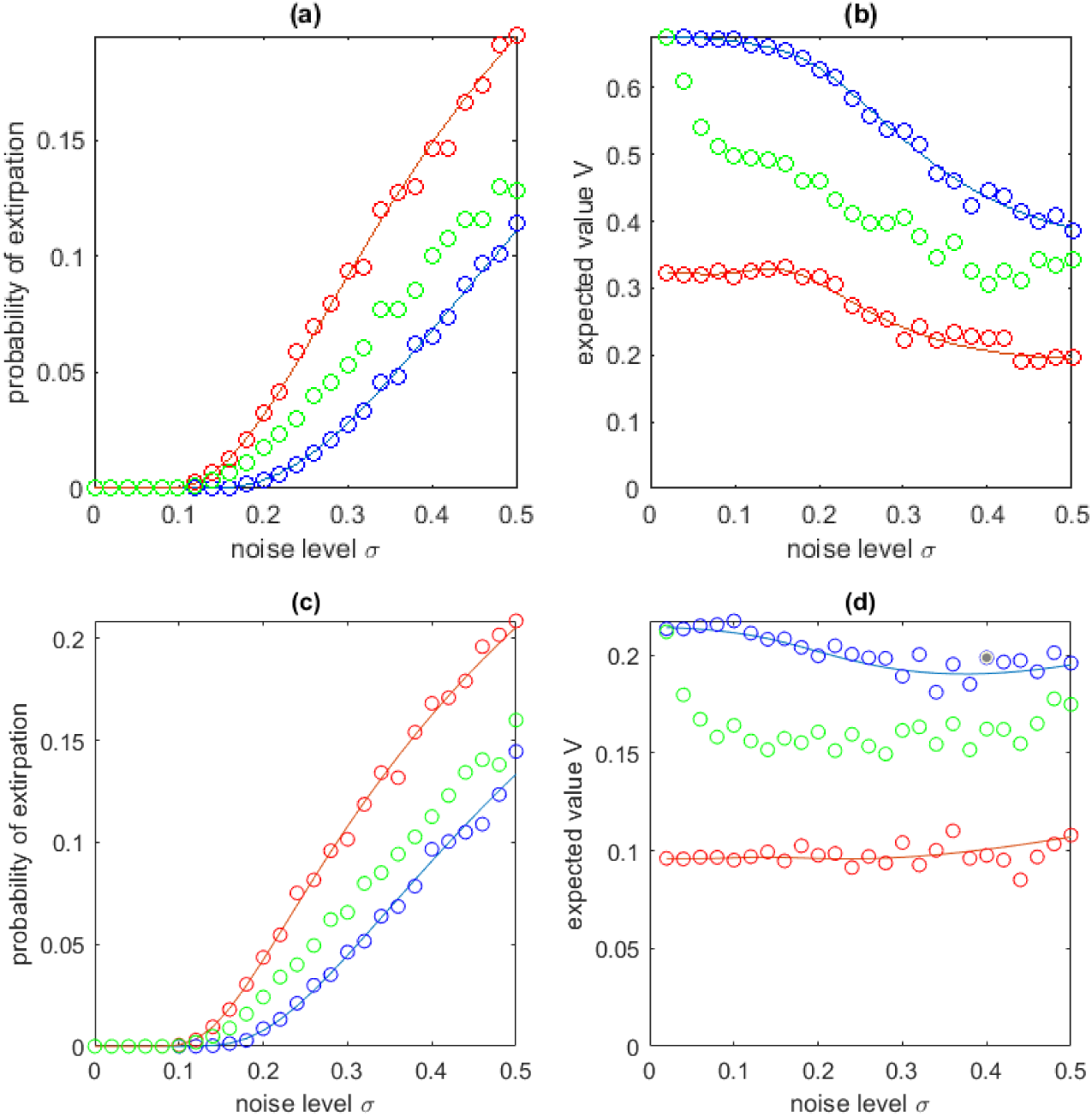
Comparison of Markov decision process solutions (solid curves) with agent-based model simulations (points): (a,c) probability of extirpation per year 1 − λ; (b,d) expected value (i.e. average long-term return from fishing). Discount factor in (a,c) *δ* = 0.9; (b,d) *δ* = 0.7. Blue curves show the results from the Markov decision process at the Pareto efficient solution 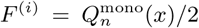; blue points show the agent-based simulation results for two agents playing ALLC. Red curves show the results from the Markov decision process under the Nash equilibrium 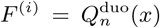; red points show the agent-based simulation results for two agents playing ALLD. Green points show the agent-based simulation results for two agents playing TFT. The agent-based model was simulated for a *n*_0_ = 20 year burn-in period which was discarded, followed by a *n*_max_ = 100 year period, and results are averaged over *M* = 500 independent realisations, discarding any realisation in which the stock is extirpated before the end of the burn-in period. Parameter values shown in Table 1

Under tit-for-tat (TFT, green), the expected value gets close to the cooperative outcome when noise is very small. However, as noise increases, the expected value rapidly drops and ends up approximately midway between the cooperative and non-cooperative outcomes. Interestingly, noise impacts the performance of TFT relative to the Pareto efficient ALLC before it affects the expected value from ALLC itself. This shows that even small amounts of noise make sustained cooperation more difficult.

The results in Figure 5(a,b) are for a discount factor of *δ* = 0.9. For comparison, we also solved the Markov decision process and simulated the agent-based model for discount factors of 0.7 and 0.95. When the discount factor is high, agents fish at slightly lower levels to protect biomass and therefore profits in future years. However, the effect of noise is similar to that seen in Fig. 5(a,b). When the discount factor is low (*δ* = 0.7, Fig. 5(c,d)), agents attach less value to profits in future years. This greatly reduces the effect of noise on the expected value, because extirpation at some future time is less important from the point of view of a short-sighted agent. However, the probability of extirpation is higher than the *δ* = 0.9 case, as a consequence of the higher average fishing mortality.

The model can also be used to simulate two agents playing different strategies. At low noise levels, the results of this are as expected from the classical iterated prisoner’s dilemma: ALLC vs ALLD results in a large advantage for the defector; ALLC vs TFT results in both agents always cooperating; ALLD vs TFT results in both agents always defecting. As noise increases, the actions of an agent playing TFT become increasingly determined by noise in the biomass signal and less sensitive to the actions of their opponent. This is the core reason why TFT ceases to be effective in the presence of noise.

## 4. Discussion

We have developed a stochastic optimisation framework for determining the optimal fishing effort for either a monopolist or duopolist operating in a noisy environment. A solution to the monopolist’s Markov decision process is a type of harvest control rule. This is an increasingly used fisheries management tool that specifies biomass reference points at which increasingly strict controls on fishing are imposed [64]. Our model assumes that fishing agents or managers can accurately observe the stock biomass once during the annual cycle of recruitment and fishing. Biomass estimates are often inaccurate due to a lack of data [21] or to measuring proxy variables such as catch per unit effort rather than biomass itself. Real fisheries have many other sources of noise operating in both the population dynamics and the harvesting process. We have studied the effect of including density-independent, normally distributed noise in a single model component, the annual stock-recruitment relationship. Our model provides a natural framework for including other sources of variability, such as uncertainty in biomass observations [64], noise in fishing mortality due to spatial heterogeneity or variations in catchability, or density-dependent noise [19]. Nevertheless, considering noise in annual recruitment is a reasonable starting point as this is the largest single source of unexplained variability in most fish stocks [13, 23, 59].

In both the monopolist and duopolist case, our results show that there is a threshold stock biomass below which it is optimal to stop fishing until the stock recovers. This threshold is lower for a duopolist than for a monopolist. Correspondingly, the stock has a lower equilibrium average biomass and a higher probability of extirpation under a duopolist than under a monopolist. The duopolist solution is optimal in the sense of being a Nash equilibrium, that is neither player can increase their expected profit by unilaterally changing their fishing effort. However, this solution is not Pareto efficient, meaning that both players could obtain a higher profit by cooperatively reducing their fishing effort. Thus the duopolist’s decision problem contains the structure of an iterated prisoner’s dilemma.

We used an agent-based model to test the effectiveness of a simple, strategy based on punishment of a player who defects from a cooperative solution, also known as a threat policy [27]. Our results quantify the likelihood of punishment being triggered by environmental noise and the longterm effect on fisheries-level outcomes. Although this strategy is effective in maintaining mutual cooperation when noise is very small, its performance rapidly drops as the noise level increases and swamps the player’s ability to discern any accurate information about their opponent’s actions. This shows that, unless players are able to make enforceable agreements about fishing effort, sustaining cooperation in a noisy environment is difficult.

Other punishment-based strategies exist, such as a grim trigger, where a single perceived defection by an opponent triggers the player to defect for the remainder of the process, or generous tit-for-tat, which has some probability of forgiving a perceived defection. The grim trigger is a subgame perfect Nash equilibrium for some iterated games, such as the infinitely iterated prisoner’s dilemma [65]. However, we investigated tit-for-tat in order to focus on a simple strategy that punishes attempts at cheating, but allows for a return to mutual cooperation following perceived defections. We do not claim that tit-for-tat is an optimal strategy: more complex strategies could also be considered that have memory of more than one round, or have a more nuanced punishment regime than just a binary choice between the Nash equilibrium and Pareto efficient fishing efforts. These could be simulated using our agent-based framework or tackled using a reinforcement learning approach [49], but we leave this for future work.

Our finding that sustained cooperation of autonomous fishing agents is difficult is an example of Hardin’s tragedy of the commons [66]. This appears to be an argument in favour of centralised controls on fishing mortality, and there are certainly many cases where these are necessary to prevent overfishing [67]. However, there are also documented cases where community management has been effective [68], particularly in small-scale fisheries [69]. We acknowledge that game theoretic models that assume selfish economic rationality do not always capture the nuances of human behaviour [70, 71].

Our models assumes that players have symmetric costs, prices and information for players. Relaxing some of these assumptions would be an interesting extension would to the study. In many asymmetric situations, side-payments become part of the optimal policy, for example an agent with a higher price to cost ratio will pay another agent to reduce their fishing effort [25, 26]. Other potential model extensions include a multi-species fishery, harvesting by multiple agents, and age- or size-structured populations.

## Acknowledgements

MH was supported by a PhD scholarship funded jointly by the University of Canterbury and Te Pūnaha Matatini. AJ and MJP were partly supported by Te Pūnaha Matatini. The authors are grateful to Dr Suzi Kerr and Dr Tava Olsen for discussions about the research and to Dr Mark Fackrell, Dr Mark McGuinness and two anonymous reviewers for comments on an earlier version of the manuscript.

## Appendix A. Solution of the Bellman equation

The Bellman equation takes the form

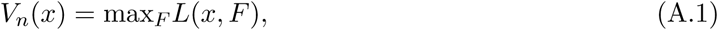

where, for the monopolist problem, using Eq. (7) gives

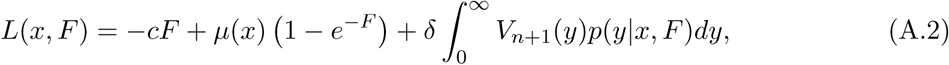

where *µ*(*x*) is the expected value of the stochastic Beverton–Holt map defined by Eq. (2), given by

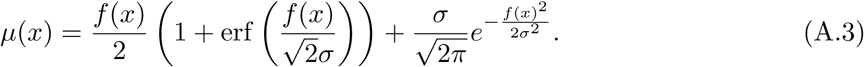

Note that if the variable *x* were unconstrained, the expected value of the stochastic map would just be the deterministic Beverton–Holt map *f* (*x*). However, because Eq. (2) constrains *x* to take non-negative values, *µ*(*x*) is greater than *f* (*x*).

Differentiating *L*(*x, F*) with respect to the action variable *F* gives

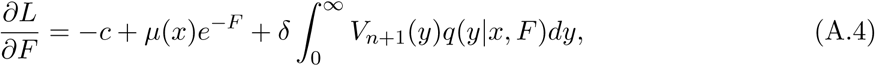

where

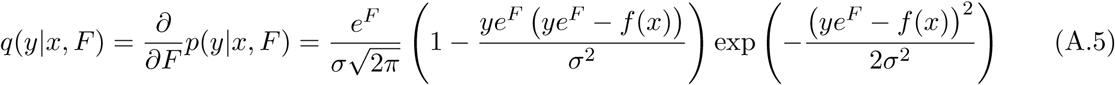

It can be shown that *∂*^2^*L/∂F* ^2^ < 0. Hence, for a given *x*, there are two possibilities: (i) if lim_*F* →0_ *∂L/∂F* > 0 then the optimal fishing mortality is the unique positive root of *∂L/∂F* = 0; (ii) if lim_*F* →0_ *∂L/∂F* < 0 then the optimal fishing mortality is *F* = 0.

The same approach can be taken to the multi-agent version of the problem, where each agent can have their own cost relative to price parameter *c*_*i*_ and discount factor *δ*_*i*_. In the general case with *m* agents, the Bellman equation takes the form

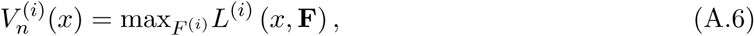

where **F** = [*F* ^(1)^, …, *F* ^(*m*)^] and

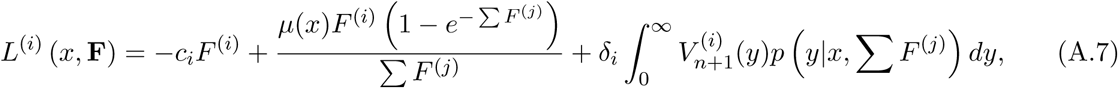

and all summations are over *j* = 1, …, *m* unless otherwise specified.

Differentiating *L*^(*i*)^ with respect to the action variable *F* ^(*i*)^ gives

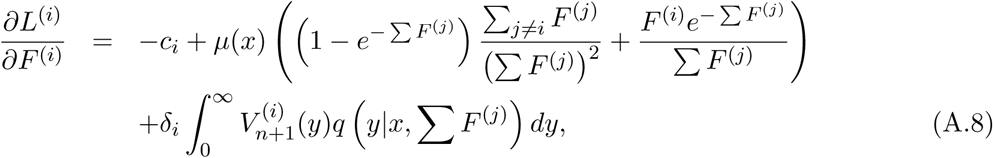

where *q*(*y*|*x, F*) is given by Eq. (A.5) as previously.

In the case of identical agents (*c*_*i*_ = *c*_*j*_, *δ*_*i*_ = *δ*_*j*_ for all *i, j*), the optimal policy must be the same for each agent *i* by symmetry. Therefore we require a solution of *∂L*^(*i*)^*/∂F* ^(*i*)^ = 0 satisfying *F* ^(*i*)^ = *F* ^(*j*)^ = *F* for all *i, j*. Thus the optimal fishing mortality for an agent when the biomass is *x* in year *n* is given by the root *F* of *M* (*x, F*) = 0 where:

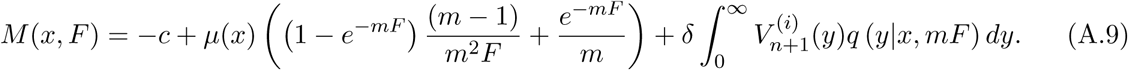

It can be shown that *∂M/∂F* < 0, so again there are two possibilities: (i) if lim_*F* →0_ *M* (*x, F*) > 0 then the optimal fishing mortality is the unique positive root of Eq. (A.9); (ii) if lim_*F* →0_ *M* (*x, F*) < 0 then the optimal fishing mortality is *F* = 0. The symmetric duopolist’s optimal policy is found by setting *m* = 2.

In the case of non-identical agents, a solution **F** is required to satisfy

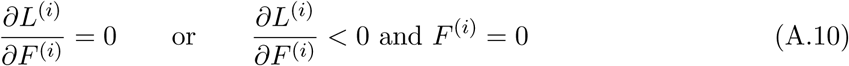

### Numerical methods

Solutions of the Bellman equation are calculated numerically by truncating the state variable *x* at *x* = *x*_max_ = 2 and discretising with step size *δx* = 0.0025.

For the monopolist’s problem and symmetric duopolist’s problem, in year *n* and state *x*_*k*_, the value of lim_*F* →0_ *M* (*x*_*k*_, *F*) is calculated. If this negative, then the optimal policy *Q*_*n*_(*x*_*k*_) is 0, otherwise *Q*_*n*_(*x*_*k*_) is the unique positive root of Eq. (A.9), which is found using Matlab’s single-variable root-finding routing *fzero*.

For the asymmetric duopolist’s problem, the optimal policy in year *n* and state *x*_*k*_ is found via an iterative procedure, using the solution for the previos state as the initial condition. This procedure calculates the optimal response of agent *i* to a fixed policy for agent *j* by solving Eq. (A.10), then the optimal response of agent *j* to agent *i* and so on, iterating until convergence.

Once the optimal policy is known, the value function *V*_*n*_(*x*_*k*_) is calculated according to Eq. (A.7) with 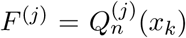 for all agents *j*. This means that the value function is tabulated at *x* = *x*_*k*_ and so the integrals in Eqs. (A.7) and (A.9) are evaluated using the trapezium rule with step size *δx*.

The value of *x*_max_ = 2 is sufficiently large that, for the noise levels considered, there is a very small probability that the biomass exceeds this. Reducing the step size to *δx* = 0.001 had a negligible effect on the results.

## References

[1] J. Hjort, Fluctuations in the great fisheries of northern Europe viewed in the light of biological research, ICES, 1914.

[2] P. J. Doherty, D. M. Williams, The replenishment of coral reef fish populations, Oceanogr Mar Biol Annu Rev 26 (48) (1988) 551.

[3] P. Lehodey, J. Alheit, M. Barange, T. Baumgartner, G. Beaugrand, K. Drinkwater, J.-M. Fromentin, S. Hare, G. Ottersen, R. Perry, et al., Climate variability, fish, and fisheries, Journal of Climate 19 (20) (2006) 5009–5030.

[4] C. M. O’Brien, C. J. Fox, B. Planque, J. Casey, Fisheries: climate variability and North Sea cod, Nature 404 (6774) (2000) 142.

[5] J. J. Pritt, E. F. Roseman, T. P. O’brien, Mechanisms driving recruitment variability in fish: comparisons between the Laurentian Great Lakes and marine systems, ICES Journal of Marine Science 71 (8) (2014) 2252–2267.

[6] S. E. Swearer, J. E. Caselle, D. W. Lea, R. R. Warner, Larval retention and recruitment in an island population of a coral-reef fish, Nature 402 (6763) (1999) 799.

[7] J. A. Wiens, Population responses to patchy environments, Annual Review of Ecology and Systematics 7 (1) (1976) 81–120.

[8] S. A. Levin, Population dynamic models in heterogeneous environments, Annual Review of Ecology and Systematics 7 (1) (1976) 287–310.

[9] A. Lomnicki, Individual differences between animals and the natural regulation of their numbers, The Journal of Animal Ecology (1978) 461–475.

[10] J. A. Tyler, K. A. Rose, Individual variability and spatial heterogeneity in fish population models, Reviews in Fish Biology and Fisheries 4 (1) (1994) 91–123.

[11] E. Houde, R. Hoyt, Fish early life dynamics and recruitment variability, Trans. Am. Fish. Soc

[12] P. Pepin, R. A. Myers, Significance of egg and larval size to recruitment variability of temperate marine fish, Canadian Journal of Fisheries and Aquatic Sciences 48 (10) (1991) 1820–1828.

[13] M. J. Fogarty, M. P. Sissenwine, E. B. Cohen, Recruitment variability and the dynamics of exploited marine populations, Trends in Ecology & Evolution 6 (8) (1991) 241–246.

[14] M. L. Puterman, Markov Decision Processes: Discrete Stochastic Dynamic Programming, John Wiley & Sons, 2014.

[15] D. J. White, Real applications of Markov decision processes, Interfaces 15 (6) (1985) 73–83.

[16] D. J. White, A survey of applications of Markov decision processes, Journal of the Operational Research Society 44 (11) (1993) 1073–1096.

[17] R. Mendelssohn, Using Markov decision models and related techniques for purposes other than simple optimization: analyzing the consequences of policy alternatives on the management of salmon runs, Fishery Bull. 78 (1) (1980) 35–50.

[18] R. Mendelssohn, Discount factors and risk aversion in managing random fish populations, Canadian Journal of Fisheries and Aquatic Sciences 39 (9) (1982) 1252–1257.

[19] R. Mendelssohn, Optimal harvesting strategies for stochastic single-species, multiage class models, Mathematical Biosciences 41 (3-4) (1978) 159–174.

[20] S. H. Mann, A mathematical theory for the harvest of natural animal populations when birth rates are dependent on total population size, Mathematical Biosciences 7 (1-2) (1970) 97–110.

[21] M. Sobel, Stochastic fishery games with myopic equilibria, Essays in the Economics of Renewable Resources (1982) 259–268.

[22] D. E. Lane, A partially observable model of decision making by fishermen, Operations Research 37 (2) (1989) 240–254.

[23] P. Sparre, S. C. Venema, Introduction to tropical fish stock assessment. Part 1. Manual, FAO Fisheries Technical Paper 306.1, Rev. 2, Food and Agriculture Organisation, Rome, 1998.

[24] M. Bailey, U. R. Sumaila, M. Lindroos, Application of game theory to fisheries over three decades, Fisheries Research 102 (1) (2010) 1–8.

[25] R. McKelvey, Game-Theoretic Insights into the International Management of Fisheries, Natural Resource Modeling 10 (2) (1997) 129–171.

[26] G. R. Munro, The optimal management of transboundary renewable resources, Canadian Journal of Economics (1979) 355–376.

[27] V. Kaitala, M. Pohjola, Optimal recovery of a shared resource stock: a differential game model with efficient memory equilibria, Natural Resource Modeling 3 (1) (1988) 91–119.

[28] C. Chiarella, M. C. Kemp, N. Van Long, K. Okuguchi, On the economics of international fisheries, in: Contributions to Economic Analysis, vol. 150, Elsevier, 189–198, 1984.

[29] U. R. Sumaila, A review of game-theoretic models of fishing, Marine Policy 23 (1) (1999) 1–10.

[30] G. Martin-Herran, J. P. Rincón-Zapatero, Efficient Markov perfect Nash equilibria: theory and application to dynamic fishery games, Journal of Economic Dynamics and Control 29 (6) (2005) 1073–1096.

[31] M. J. Plank, J. Kolding, R. Law, H. D. Gerritsen, D. Reid, Balanced harvesting can emerge from fishing decisions by individual fishers in a small-scale fishery, Fish and Fisheries.

[32] M. Hackney, A. James, M. J. Plank, Emergence of balanced harvesting in an agent-based model of an open-access small-scale fishery, Mathematical biosciences 316 (2019) 108245.

[33] S. D. Fretwell, H. L. Lucas Jr., On territorial behavior and other factors influencing habitat distribution in birds. I. Theoretical development, Acta Biotheoretica 19 (1969) 16–36.

[34] J. M. Smith, Evolution and the Theory of Games, Cambridge university press, 1982.

[35] K. Regelmann, Competitive resource sharing: a simulation model, Animal Behaviour 32 (1) (1984) 226–232.

[36] A. Kacelnik, J. R. Krebs, C. Bernstein, The ideal free distribution and predator-prey populations, Trends in Ecology and Evolution 7 (2) (1992) 50–55.

[37] D. M. Gillis, R. M. Peterman, A. V. Tyler, Movement dynamics in a fishery: application of the ideal free distribution to spatial allocation of effort, Canadian Journal of Fisheries and Aquatic Sciences 50 (2) (1993) 323–333.

[38] D. M. Gillis, A. van der Lee, Advancing the application of the ideal free distribution to spatial models of fishing effort: the isodar approach, Canadian Journal of Fisheries and Aquatic Sciences 69 (10) (2012) 1610–1620.

[39] I. E. van Putten, S. Kulmala, O. Thébaud, N. Dowling, K. G. Hamon, T. Hutton, S. Pascoe, Theories and behavioural drivers underlying fleet dynamics models, Fish and Fisheries 13 (2) (2012) 216–235.

[40] M. J. Sobel, Myopic solutions of Markov decision processes and stochastic games, Operations Research 29 (5) (1981) 995–1009.

[41] M. L. Littman, Markov games as a framework for multi-agent reinforcement learning, in: Machine Learning Proceedings 1994, Elsevier, 157–163, 1994.

[42] M. Bowling, M. Veloso, An analysis of stochastic game theory for multiagent reinforcement learning, Tech. Rep., Carnegie-Mellon Univ, Pittsburgh P.A., School of Computer Science, 2000.

[43] M. Zinkevich, A. Greenwald, M. L. Littman, Cyclic equilibria in Markov games, in: Advances in Neural Information Processing Systems, 1641–1648, 2006.

[44] J. Filar, K. Vrieze, Competitive Markov Decision Processes, Springer Science & Business Media, 2012.

[45] O. J. Vrieze, Stochastic games with finite state and action spaces, CWI tracts.

[46] J. Hu, M. P. Wellman, Nash Q-learning for general-sum stochastic games, Journal of machine learning research 4 (Nov) (2003) 1039–1069.

[47] J. Hu, M. P. Wellman, et al., Multiagent reinforcement learning: theoretical framework and an algorithm., in: ICML, vol. 98, Citeseer, 242–250, 1998.

[48] M. L. Littman, Value-function reinforcement learning in Markov games, Cognitive Systems Research 2 (1) (2001) 55–66.

[49] R. S. Sutton, A. G. Barto, Reinforcement Learning: An Introduction, MIT press, 2018.

[50] W. B. Powell, Approximate Dynamic Programming: Solving the Curses of Dimensionality, vol. 703, John Wiley & Sons, 2007.

[51] J. Kober, J. A. Bagnell, J. Peters, Reinforcement learning in robotics: A survey, The International Journal of Robotics Research 32 (11) (2013) 1238–1274.

[52] Y. Niv, Reinforcement learning in the brain, Journal of Mathematical Psychology 53 (3) (2009) 139–154.

[53] E. Ponomarev, I. Oseledets, A. Cichocki, Using Reinforcement Learning in the Algorithmic Trading Problem, Journal of Communications Technology and Electronics 64 (12) (2019) 1450–1457.

[54] D. Fudenberg, J. Tirole, Game Theory, Cambridge, Massachusetts. MIT press, 1991.

[55] D. Booth, G. Beretta, Seasonal recruitment, habitat associations and survival of pomacentrid reef fish in the US Virgin Islands, Coral Reefs 13 (2) (1994) 81–89.

[56] D. M. Allen, D. L. Barker, Interannual variations in larval fish recruitment to estuarine epibenthic habitats., Marine Ecology Progress Series. Oldendorf 63 (2) (1990) 113–125.

[57] S. E. Campana, Year-class strength and growth rate in young Atlantic cod Gadus morhua, Marine Ecology Progress Series 135 (1996) 21–26.

[58] D. O. Conover, T. M. Present, Countergradient variation in growth rate: compensation for length of the growing season among Atlantic silversides from different latitudes, Oecologia 83 (3) (1990) 316–324.

[59] R. J. Beverton, S. J. Holt, On the dynamics of exploited fish populations, vol. 11, Springer Science & Business Media, 1957/2012.

[60] W. E. Ricker, Stock and recruitment, Journal of the Fisheries Board of Canada 11 (5) (1954) 559–623.

[61] R. Bellman, et al., The theory of dynamic programming, Bulletin of the American Mathematical Society 60 (6) (1954) 503–515.

[62] R. Bellman, Dynamic programming, Science 153 (3731) (1966) 34–37.

[63] R. Axelrod, The evolution of strategies in the iterated prisoners dilemma, The Dynamics of Norms (1987) 1–16.

[64] S. F. Kvamsdal, A. Eide, N.-A. Ekerhovd, K. Enberg, A. Gudmundsdottir, A. H. Hoel, K. E. Mills, F. J. Mueter, L. Ravn-Jonsen, L. K. Sandal, et al., Harvest control rules in modern fisheries management, Elementa Science of the Anthropocene 4 (2016) 000114.

[65] S. Tadelis, Game Theory: An Introduction, Princeton University Press, 2013.

[66] G. Hardin, The Tragedy of the Commons, Science 162 (3859) (1968) 1243–1248.

[67] R. Hilborn, D. Ovando, Reflections on the success of traditional fisheries management, ICES journal of Marine Science 71 (5) (2014) 1040–1046.

[68] E. Ostrom, Governing the commons: The evolution of institutions for collective action, Cambridge University Press, Cambridge, 1990.

[69] J. Kolding, P. A. M. van Zwieten, The tragedy of our legacy: How do global management discourses affect small scale fisheries in the South?, Forum for Development Studies 38 (2011) 267–297.

[70] E. A. Fulton, A. D. M. Smith, D. C. Smith, I. E. van Putten, Human behaviour: the key source of uncertainty in fisheries management, Fish and fisheries 12 (1) (2011) 2–17.

[71] A. Mashanova, R. Law, Resource dynamics, social interactions, and the tragedy of the commons, in: H. Liljenstrom, U. Svedin (Eds.), Micro, Meso, Macro: Addressing Complex Systems Couplings, World Scientific, Singapore, 171–183, 2005.

